# Shock drives a highly coordinated transcriptional and DNA methylation response in the endothelium

**DOI:** 10.1101/2022.08.01.502384

**Authors:** Ramon Bossardi Ramos, Nina Martino, Dareen Chuy, Shuhan Lu, Uma Balasubramanian, Iria Di John Portela, Peter A. Vincent, Alejandro P. Adam

## Abstract

Endothelial dysfunction is a critical factor in promoting organ failure during septic shock. Organ dysfunction during shock increases the risk of long-term sequelae in survivors through mechanisms that remain unknown. We postulated that vascular dysfunction during shock contributes to the long-term morbidity post shock through transcriptional and epigenetic changes within the endothelium. As we have previously demonstrated that IL-6/JAK/STAT3 signaling in endothelial cells contributes to the inflammatory response following endotoxin, we performed cross-omics analyses on kidney endothelium from acute endotoxin-challenged mice lacking or not the JAK/STAT3 inhibitor SOCS3. This analysis revealed significant DNA methylation changes upon proinflammatory signaling that was significantly associated with transcriptional activity through AP1, STAT, and IRF families, suggesting a mechanism driving transcription-induced gene-specific methylation changes. In vitro, we demonstrated that IL-6 induces similar changes in DNA methylation. Specific genes showed DNA methylation changes in response to an IL-6+R challenge, and consistently, changes in their expression levels by 72 hours of IL-6+R treatment. Further, changes in the endothelial methylome remain in place for prolonged periods in absence of IL-6, suggesting that this cytokine may elicit transcriptional changes long after the resolution of inflammation. Also, demonstrated that DNA methylation changes could directly alter the expression of these genes and that STAT3 activation had a causal role in this transcriptional response. Our findings provide evidence for a critical role for IL-6 signaling in regulating shock-induced epigenetic changes and sustained endothelial activation, offering a new therapeutic target to limit vascular dysfunction and prevent long-term organ damage.

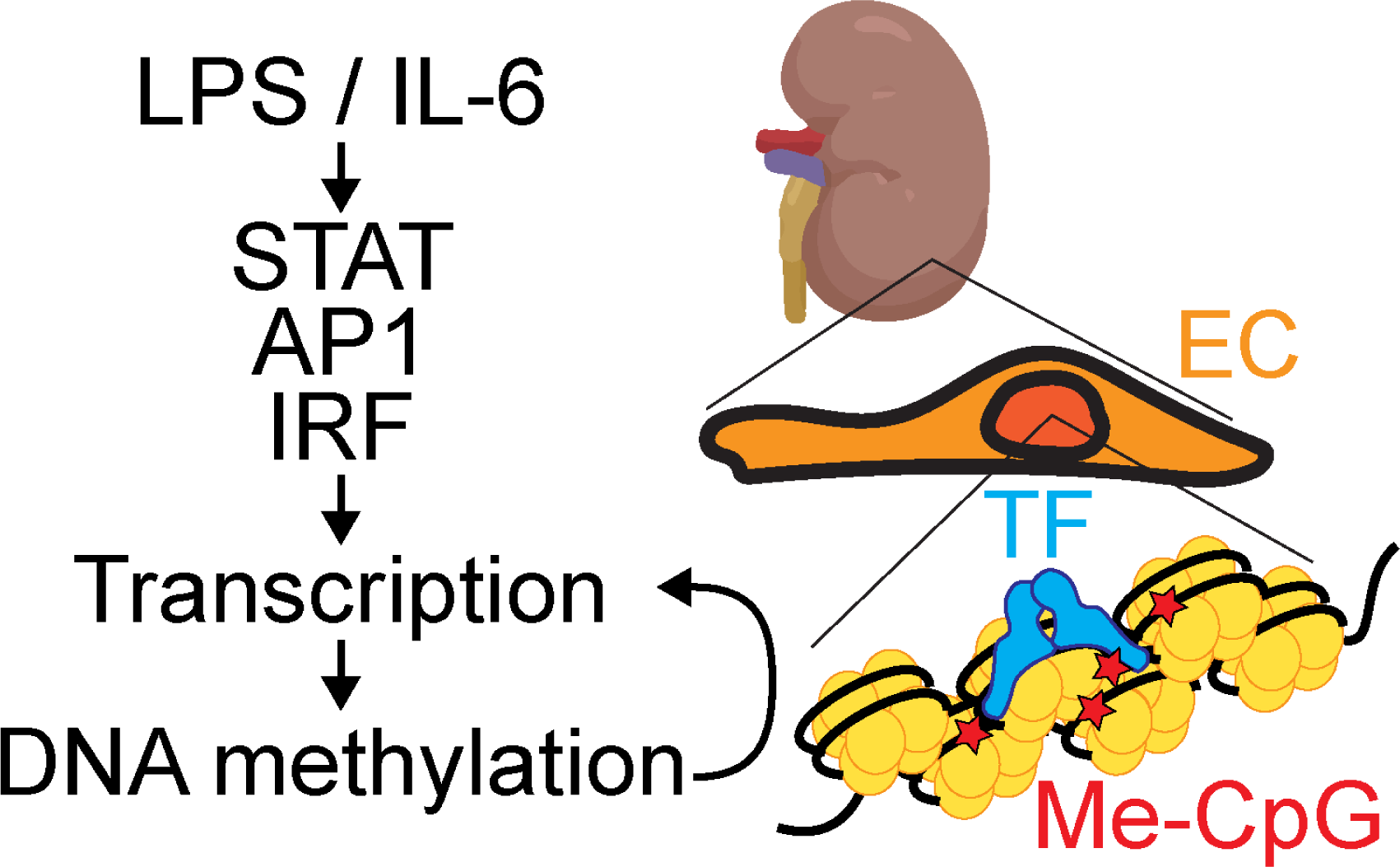

## Introduction

Shock-induced endothelial dysfunction is a critical factor leading to organ failure^1^ by promoting vascular leak, leukocyte adhesion, thrombosis and loss of vasoreactivity^2^. While advances in critical care medicine resulted in an increased survival rate for subjects with critical illness^3–5^, survivors continue to succumb to sequelae months and even years after the initial acute episode, a condition known as post-intensive care syndrome (PICS). This syndrome involves long-term physical, cognitive and mental deficits and carries increased 5-year mortality rates^3, 6–8^. The mechanisms driving PICS are not understood. Extensive efforts have been made to define the role of epigenetic regulation, including DNA methylation, in sepsis.

Shock is associated with an acute release of cytokines, a process called cytokine storm^7^. Among all the cytokines changed during a cytokine storm, the circulating levels of IL-6 are particularly informative, as they correlate with systemic disease severity^9, 10^ and are highly predictive of mortality^11^. Moreover, IL-6 signaling induces its own expression through a positive feedback loop that is associated with worse outcomes^9, 10, 12^. The effects of this cytokine in the endothelium are mediated by IL-6 trans-signaling^13^, which involves a ternary binding between the cytokine, a circulating soluble form of the IL-6-binding receptor subunit (sIL6Rα), and the transmembrane receptor subunit gp130 to induce JAK kinase activity and STAT3 phosphorylation. This pathway is inhibited by a negative feedback loop involving de novo synthesis of SOCS3^9, 10^. Previous work from our lab demonstrated that IL-6 signaling promotes STAT3-dependent sustained barrier function loss in human endothelial cells (EC) primary cultures^14^, and we recently showed in primary ECs that this signaling axis functions as a critical autocrine loop induced by endotoxin (lipopolysaccharides from *E Coli*, LPS) or TNF^15^. Further, we demonstrated that endothelial-specific, tamoxifen-inducible SOCS3 knockout (SOCS3iEKO) mice display mortality as soon as 18 hours after an endotoxin challenge, which is preceded by increased IL-6 endothelial levels and severe vasculopathy leading to kidney failure^12^. The organ damage was associated with a kidney proinflammatory transcriptional profile consistent with increased leukocyte adhesivity and thrombogenicity of the endothelial surface, including high levels of COX2, VWF and P-selectin^12^. The specific mechanisms driving SOCS3-dependent endothelial dysfunction in this context, however, remains unknown.

Circulating cells in septic patients show substantial DNA methylation changes^16–18^, which have been suggested to contribute to immune activation and tolerance^19, 20^. It is conceivable that similar epigenetic changes occur within the affected organs, but limitations in the availability of septic biopsies greatly restricts our understanding beyond circulating cells. Despite its clinical implications, the endothelial epigenetic changes and the transcription factors (TF) network that mediate endothelial gene expression changes in response to shock are not fully understood. A better understanding of the epigenetic, TF and transcriptional changes driving endothelial dysfunction may identify novel biomarkers to predict or provide potent therapeutic targets to treat long-term endothelial disfunction.

In this study, we sought to determine the transcriptional and epigenetic changes that occur in endothelial cells in response to proinflammatory stimuli initiated by endotoxin. We performed cross-omics analyses of transcriptome and DNA methylome data obtained using DNA collected from isolated kidney endothelium of wild type (WT) and SOCS3iEKO mice challenged with endotoxin, and from human umbilical vein endothelial cells (HUVEC) treated with IL-6. This analysis allowed us to obtain a comprehensive assessment of gene regulatory relationships. In addition, we performed motif enrichment and discriminant regulon expression analysis to identify the transcription factors associated with these responses. We found that a single endotoxin challenge leads to substantial DNA methylation changes in the kidney endothelium and SOCS3iEKO promotes a divergent response in DNA methylation in following exposure to LPS. Further, we found that increased STAT and AP1 (JunB, BATF, and cFos) transcriptional activity was associated with DNA methylation changes, suggesting a potential mechanism driving transcription-induced gene-specific methylation changes. In vitro, we demonstrated that IL-6 induces similar changes in DNA methylation and that many changes in the endothelial methylome remain in place even without continued IL-6 signaling. Motif enrichment analysis suggested that not only STAT3, but also JunB, BATF, and cFos were associated with both epigenetic and transcriptional changes. Depletion of STAT3 and JunB using siRNA in HUVEC demonstrated a role for these two TFs in many of these changes. Together, these findings demonstrate that the endothelium responds to shock with transcriptional and DNA methylation changes, at least in part via the coordination of multiple transcription factors, including STAT1, STAT3 and the AP1 family members cFos, JunB and BATF.

## Material and Methods

The commercial sources for critical reagents and their catalog numbers are listed in Supplemental Table 9. Supplemental Table 10 provides a list of sequences for RT-qPCR primers. Supplemental Table 11 lists all antibodies used.

### Mice

All animal experiments were approved and conducted in accordance with the Albany Medical College IACUC guidelines for animal care and were performed in the Animal Research Facility at Albany Medical College. Mice were housed in specific pathogen-free rooms with 12-hour light/12-hour dark cycle and controlled temperature and humidity. Mice were kept in groups of 5 or fewer in Allentown cages with access to food and water ad libitum. Endothelial-specific, tamoxifen-inducible SOCS3 knockout (SOCS3iEKO) mice on a C57Bl6/J background were described previously^12^. Briefly, all mice carried two copies of a Cdh5-CreERT2 endothelial driver^21^ and ROSA26-tdTomato reporter^22^ by crossing B6.Tg(Cdh5-cre/ERT2)1Rha with B6.Gt(ROSA)26Sor^tm9^(CAG–tdTomato)^Hze^. A floxed SOCS3 transgene^23^ was introduced by breeding to B6;129S4-Socs3^tm1Ayos^/J to generate the SOCS3iEKO. Control and SOCS3iEKO littermates were obtained by breeding two SOCS3^fl/+^ heterozygous mice. Genotyping primers have been described previously^12^. All mice received tamoxifen (2 mg tamoxifen in 100 μL via intraperitoneal (IP) injection) at 6–9 weeks old for 5 consecutive days. All the experiments were conducted between 2 and 3 weeks after the end of tamoxifen treatment and the SOCS3 deletion gene was confirmed by post-tamoxifen tail digestion and PCR.

Severe, acute inflammation was induced by a single IP injection of a bolus of 250 μg/250 μl LPS. Control mice were given 250 μl of sterile saline via IP injection. Disease severity was scored 14 hours after the LPS challenge as described before^12^. Immediately after scoring, body weight and temperature were measured. Mice were then euthanized with an overdose of pentobarbital. Assignment to the saline or LPS groups was performed through randomization of mice within each genotype and sex for each litter. All handling, measurements, and scoring were performed by a researcher masked to treatment and genotype groups and based on mouse ID (ear tags). Experimental groups were unmasked at the end of each experiment.

### Enrichment of endothelial cells

After euthanasia, the animals were cannulated on the left ventricle, and the right atrium was nicked to allow perfusion with 5 ml/min PBS for 3 min. Kidneys were collected and minced into small pieces (∼1 mm^3^). The tissue was transferred to 50 ml conical tubes with 5 ml of PBS containing calcium and magnesium plus 100 mg of type 1 collagenase and 100 mg dispase II for 10 minutes at 37°C. The slurry of tissue and buffer was triturated by passage through a 14-gauge cannula attached to a 10 ml syringe approximately 10 times, followed by a 20-gauge cannula attached to a 10 ml syringe. The cell suspensions were filtered through a 70 µm cell strainer, resulting in single cell suspensions prior to centrifugation at 400×g for 8 min at 4 °C. Pellets were resuspended with 1.5ml of ice-cold PBS + 0.1% BSA. One aliquot (75 μl) per sample was lysed in Trizol to obtain whole kidney RNA.

The remainder cell suspension was transferred to RNAse-free polystyrene tubes containing 60 µl of streptavidin-conjugated dynabeads bound to biotinylated EpCAM and biotinylated CD45 antibodies. The cell suspensions were incubated with these beads for 10 min at RT with rotation. Following this, tubes were placed on a Dynal MPC-S magnet for 5 minutes. The supernatants were transferred to RNAse free polystyrene tubes containing 45µl of streptavidin-conjugated dynabeads bound to biotinylated CD31 antibodies and incubated for 10 min at RT with rotation. Following this, tubes were placed on a Dynal MPC-S magnet for 5 minutes and the supernatant was discarded. The remaining cells bound to the beads were washed six times with PBS containing 0.1% BSA to remove any contaminating cells. After removal of the final wash, the cells were resuspended in 400 µl PBS. From this, 200 µl was transferred to a RNAse-free tube and DNA was isolated using the DNeasy Blood & Tissue Kit (Qiagen, USA). The other 200 µl were used for RNA isolation following the RNeasy Plus Mini Kit (Qiagen) protocol. VWF and CDH1 expression were measured by RT-qPCR. The ratios of VWF and CDH1 expression in the enriched versus the total RNA from the same organ were calculated to confirm the enrichment of the endothelial cells following this protocol.

### Cell culture

Human Umbilical Vein Endothelial Cells (HUVECs) were isolated in-house according to established protocols^24–26^ as described before^12, 14, 15^. The identity and purity of the HUVEC isolations were confirmed for each isolation by more than 99% positive immunostaining with endothelial cell markers (FITC-Ulex europaeus lectin, VE-cadherin) and more than 99.9% negative for α–smooth muscle actin. Cells were assayed between passages 3 and 8. To induce IL-6 signaling, cells were plated at full confluence at a density of 8 × 10^4^ cells/cm^2^ on plates precoated for 30 min with 0.1% gelatin and incubated at least 48 h prior to the start of experiments. Cells were then treated with a combination of 200 ng/mL recombinant human IL-6 and 100 ng/mL sIL-6Rα (IL-6+R) or PBS (control) for 72 hours. After 72 hours, a subset of control and IL-6+R-treated cells were washed and incubated for 96 hours in EGM-2 Growth Medium. Treated HUVECs and controls were immediately lysed in Trizol or centrifuged and then stored until DNA extraction.

### 5-Aza-2′-deoxycytidine (5-AZA) treatment

HUVEC were seeded in six-well plate and exposed to 5-AZA (Sigma-Aldrich) at a concentration of 5μM for 72h; new 5-AZA was added each 24 hours. The control group was treated in parallel with PBS. RNA was harvested for downstream analysis.

### Immunofluorescence Microscopy

Immunofluorescence studies were performed by seeding 8x 10^4^ cells/well on 8-well μ-slide chambers (Ibidi) precoated with 0.1% gelatin. At 48 h after seeding, cells were treated as indicated. Cells were then fixed with 4% paraformaldehyde (Affymetrix) in PBS for 30 min at 4C, washed twice with PBS, and processed for immunofluorescence at room temperature. Briefly, cells were permeabilized with 0.1% Triton X-100 (Sigma) in PBS (PBS-TX) for 15 min, and blocked with 5% bovine serum in PBS-TX for 1 h. Antibodies were incubated for 2 h at room temperature. Slides were then washed in PBS-TX and stained with Alexa Fluor–conjugated secondary antibodies and 0.5 μg/mL DAPI for 1 hour at RT. Slides were then washed in PBS. Images were taken at magnification 20x and 63x using a Zeiss AXIO Observer Z1 microscope.

### RT-qPCR

Cells grown on multi-well plates or minced mice tissue was digested to single cell suspension and Ab selection were lysed with Trizol reagent (Thermo Fisher), RNA was extracted with chloroform and precipitated with isopropanol per manufacturer’s instructions. A total of 400 ng of RNA was used to prepare cDNA using Primescript RT Master Mix (Clontech) at 42 °C following manufacturer’s instructions. The cDNA was diluted 10-fold in nuclease-free water. Then, 2 μL of cDNA was used per PCR reaction. qPCR was performed in a StepOnePlus (Applied Biosystems) instrument using 10 μL SYBR green-based iTaq supermix (Bio-Rad), 7.8 μL of water, and 2 pmol primers (0.2 μL) (Thermo Fisher). Fold induction was calculated via the ΔΔCt method using GAPDH (human) or HPRT (mice) as the housekeeping gene.

### DNA isolation, bisulfite treatment, and methylation profiling

DNA was isolated using the DNeasy Blood & Tissue Kit (Qiagen, USA) according to the manufacturer’s instructions, including optional treatment with 100 mg/ml RNase A. The concentration of DNA was measured by the Qubit dsDNA BR Assay Kit (Molecular Probes). The DNA samples were stored at -80°C. Bisulfite conversion of 500 ng of genomic DNA was performed using an EZ DNA Methylation-Gold™ Kit (Zymo Research) following manufacturer’s instructions. Analysis of DNA methylation was carried out using Illumina Infinium MethylationEPIC BeadChip arrays^27^ for human genomes (>850K sites) and Infinium Mouse Methylation BeadChip arrays for mouse genomes (>285K sites) by the Epigenomic Services from Diagenode.

### DNA methylation data processing

The methylation data discussed in this publication have been deposited in NCBI’s Gene Expression Omnibus and are accessible through GEO Series accession number GSE223381 for HUVEC and GSE223336 for mice data. Methylation array data were processed with the statistical language R (version 4.2.2) and Bioconductor packages following workflows outlined in the supplemental methods. Briefly, processing of the raw HUVEC methylation data was performed with the Chip Analysis Methylation Pipeline (ChAMP) package (version 2.21.1)^28^. Raw methylation data were imported by the minfi method^29, 30^. Probes with a detection p-value >0.01 in one or more samples, probes with a bead count <3 in at least 5% of samples, non-CpG probes, probes that align to multiple locations, and sex chromosome-specific probes were removed from subsequent analyses. As a result, the remaining 727,127 probes were utilized for data analysis. Data normalization was performed by the BMIQ method^31^. Batch effect prediction was made by singular value decomposition (SVD)^32^. Mouse probe, and intensity data for each sample were imported into R using the package RnBeads (version 2.10.0)^33^, followed by preprocessing to filter probes outside the CpG context, SNP probes, sex chromosome probes, probes without intensity value, and probes with a low standard deviation^29, 34, 35^. The RnBeads.mm10 package was used to annotate the location of each probe. Downstream data analysis was performed using β values. The β value is the ratio of the methylated probe intensity to the overall intensity (the sum of the methylated and unmethylated probe intensities). βvalues range from 0 to 1, in which 0 is no methylation and 1 is complete methylation and were used to derive heatmaps and further analysis^36^. Differential methylation between groups was defined as Δβ value difference and adjusted p-value (Benjamini-Hochberg method, FDR <0.05).

Gene ontology (GO) analysis of the differentially hypomethylated or hypermethylated CpG sites associated with gene locations was performed using Metascape (version 3.5)^37^ and selecting GO terms with a p-value less than 0.05. Motif enrichment of the same CpG sets was done using findMotifsGenome.pl program of HOMER (version 4.11)^38^ and selecting a 500-bp window upstream and downstream of the differentially methylated CpG sites. Annotated CpGs in the EPIC array were used as background.

### Bulk RNA-seq data and differential expression analysis

The bulk HUVEC RNA-seq data discussed in this publication have been deposited in NCBI’s Gene Expression Omnibus and are accessible through GEO Series accession number GSE225236. RNA-seq libraries were prepared using Illumina’s TruSeq protocol and were sequenced on an Illumina NextSeq 500. Reads were aligned to the hg38 genome using Rsubread v1.5.3^39^. Gene counts were quantified by Entrez Gene IDs using featureCounts and Rsubread’s built-in annotation^40^. Gene symbols were provided by NCBI gene annotation. Genes with count-per-million above 0.5 in at least 3 samples were kept in the analysis. Differential expression analysis was performed using limma-voom^41^.

Gene Set Enrichment Analysis (GSEA) was performed comparing HUVEC treated with IL-6 or PBS for 72 hours. As gene sets collection, hallmarks (H) from the Molecular Signatures Database (MSigDB) were selected, adding the specified custom genesets. GSEA analysis and graphs were created with the ClusterProfiler^42^ and enrichplot Bioconductor packages.

### Endothelial gene expression by TRAP/RNA-Seq

We used mice carrying a TRAP transgene for translating ribosome affinity purification assays and mediated RNA isolation from the endothelium of endotoxin treated mice. These transgenic mice have a ribosomal subunit that can be replaced by a floxed GFP-tagged version (EGFP/Rpl10a), rendering the ribosome amenable to immunoprecipitation via two specific monoclonal antibodies^43^. Using this system under the endothelial-specific, tamoxifen-inducible cdh5-CreERT2 driver^21^ we isolated mRNAs that were actively translated in the endothelium of the kidneys.

14 hours after LPS treatment, mice kidneys were isolated, minced, and lysed with Trizol reagent (Thermo Fisher), as described above. RNA quantity was determined using a Qubit Flex Fluorometer (Thermo Fisher Scientific, San Jose, CA). Library preparation was performed using an Ion Chef System (Thermo Fisher Scientific, San Jose, CA) followed by sequencing using an Ion GeneStudio S5 Plus System (Thermo Fisher Scientific, San Jose, CA) both following manufacturer’s suggested protocols for the Ion AmpliSeq Transcriptome Mouse Gene Expression Kit (Thermo Fisher Scientific, San Jose, CA). Differentially expressed genes (DEGs) were identified using the Transcriptome Analysis Console (TAC) software (version 4.0.2) with the ampliSeqRNA plugin (Thermofisher Scientific). The TRAP/RNA-Seq data has been deposited in NCBI’s Gene Expression Omnibus and are accessible through GEO Series accession number GSE229292. Metascape was performed as described before.

### Gene Knockdown

Small interference RNA (siRNA) against STAT3 and JunB oligonucleotides were obtained as a set of individual siRNAs from Horizon Discovery (On target plus SiRNA). Cells were transfected with individual siRNA complexed with Lipofectamine siRNA iMAX (Invitrogen) in suspension and seeded at 1x10^5^ cells/cm2. Controls were transfected with On target plus nontargeting control pooled duplexes (Horizon Discovery). Knockdown efficiency was determined by Western blot and qPCR analysis.

### Code availability

Code files are available from the GitHub repository https://github.com/ramonbossardi/HUVEC_methylation_geneexpression

## Results

### LPS induces altered DNA methylation and transcriptional changes in the kidney endothelium

The epigenetic program in the endothelium of failing organs is not known. To begin addressing this question, we first challenged young adult mice with a single dose of endotoxin (250 μg/mouse of lipopolysaccharides from E. coli 0111:B4, LPS) or saline solution and obtained kidney endothelial DNA for methylation assays 14 hours post endotoxin injection. We previously demonstrated that this sub-lethal treatment leads to severe but transient kidney injury in WT mice^12^. As expected, we observed that LPS induced a systemic response 14 hours post-challenge, as measured by a increase in the severity score and hypothermia (Figures 1A and 1B). We enriched kidney EC from single cell suspensions using a two-step approach, first using negative selection of non-EC using EpCAM- and CD45-labeled magnetic beads, followed by a positive selection with PECAM-labeled beads (Figure 1C). EC enrichment was confirmed by measuring the expression levels of CDH1 (an epithelial marker) and VWF (an endothelial marker) in the RNA obtained from the same kidney suspension before and after magnetic bead enrichment (Figure 1D). We then performed DNA bisulfite conversion and used bead arrays to investigate the DNA methylation status of 285,000 CpG sites across the entire mouse genome. The endothelium from endotoxemic kidneys displayed 1792 CpG sites with significantly higher methylation levels (hypermethylated) and 804 CpG sites with significantly lower methylation levels (hypomethylated) than control endothelium (Figure 1E and Supplemental Table 1). Gene ontology (GO) analysis of the differentially methylated genes showed an enrichment in expected categories such as lymphocyte activation, cytokine signaling, and cell adhesion, as well as in epigenetic regulation, such as DNA methylation itself and heterochromatin assembly (Figure 1F).

**Figure 1.**
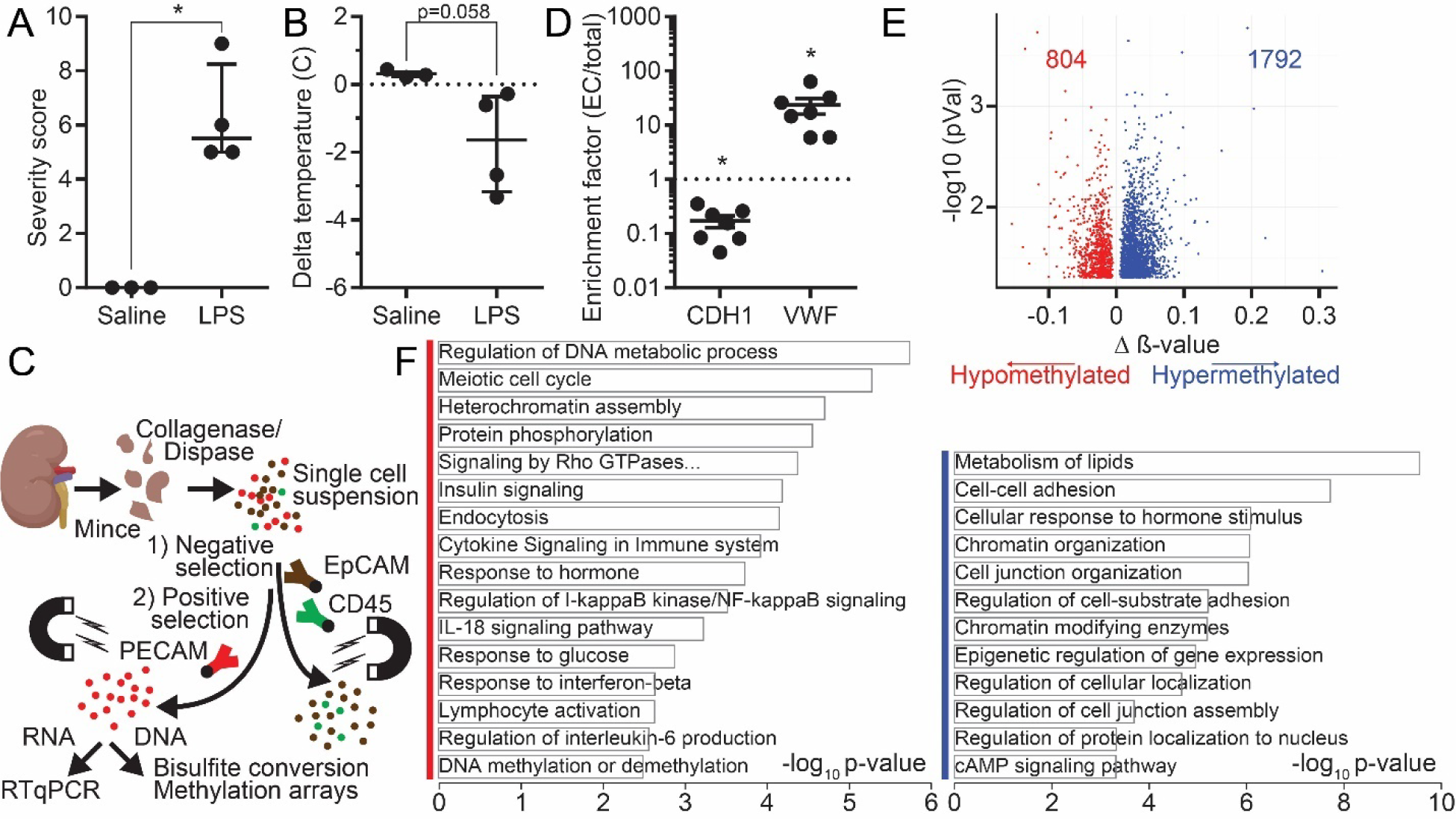
LPS-induced DNA methylation changes in the mouse kidney endothelium are enriched in genes associated with inflammatory and epigenetic responses. (**A**) Severity score of mice 14 hours after saline or LPS injection, mean ± SE. (**B**) Temperature for each mouse was measured immediately before and 14 hours post-injection, mean ± SE. (**C**) Schematic of the experimental approach for isolating kidney endothelial cells. (**D**) Expression of the epithelial marker CDH1 and the endothelial marker vWF in endothelial vs. total RNA from the same organ. (**E**) Volcano plot of differentially methylated positions CpG sites showing the p-value vs. Δβ for control and LPS-treated mice. The red dots represent significantly hypomethylated CpGs, the blue dots represent significantly hypermethylated CpGs. (**F**) Gene ontology analysis of genes associated with differentially methylated CpG sites showing the most relevant enriched categories (blue mark for the hypermethylated set, and red mark for the hypomethylated set) processed via Metascape. Asterisks denote p < 0.05.

### Loss of endothelial SOCS3 promotes a divergent response to LPS

We previously showed that SOCS3iEKO mice succumb to the same LPS challenge as above, at least in part due to kidney failure^12^. This response was associated with increased IL-6 signaling. To determine if loss of SOCS3 leads to altered DNA methylation in response to inflammatory signaling, we measured DNA methylation in the kidney endothelium in control or SOCS3iEKO mice challenged with LPS or not. Proinflammatory signaling in kidney endotoxemic endothelium of SOCS3iEKO mice was confirmed by increased expression of IL-6 and COX2 (Figure 2A). Notably, lack of endothelial SOCS3 led to a highly divergent response, with 3564 differentially methylated positions (DMP) in response to LPS in SOCS3iEKO, 2189 CpG sites that were significantly hypomethylated and 1374 sites that were significantly hypermethylated compared to the changes with control mice (Figure 2B, Supplemental Table 2). Gene ontology analyses revealed further changes in inflammation, leukocyte differentiation and cytokine signaling, as well as changes in the mitotic cell cycle, MAPK signaling, regulation of cell death, and vascular remodeling (Figure 2C). Notably, together with epigenetic regulation genes, we found multiple hypermethylated genes involved with regulation of MECP2, a 5-methylcytosine binding protein, and many hypomethylated genes associated with the transcriptional response by MECP2 (Figure 2C). To gain insight into potential roles for acute transcription and identify the transcription factors binding to regulatory elements driving these changes in endothelial DNA methylation, we performed motif enrichment analysis of these differentially methylated genes (Figure 2D). We have previously shown that loss of SOCS3 led to the expression of a type I interferon-like gene signature in failing organs^12^. Consistent with this response, hypermethylated CpG sites of LPS-treated endothelial cells in SOCS3iEKO mice were enriched in motifs binding several members of the interferon regulatory family (IRF) family. Notably, we also identified a strong enrichment in motifs binding AP1 family members JunB, ATF3, and BATF. All three genes are highly induced by IL-6 signaling in endothelial cells^12^. In contrast, hypomethylated CpGs contained motifs for several homeobox factors. Other binding sites, such as that for CTCF, were present in both groups (Figure 2D). Collectively, these results revealed specific TFs associated with both groups, suggesting a potential mechanism driving gene-specific methylation changes through transcription factor binding.

**Figure 2.**
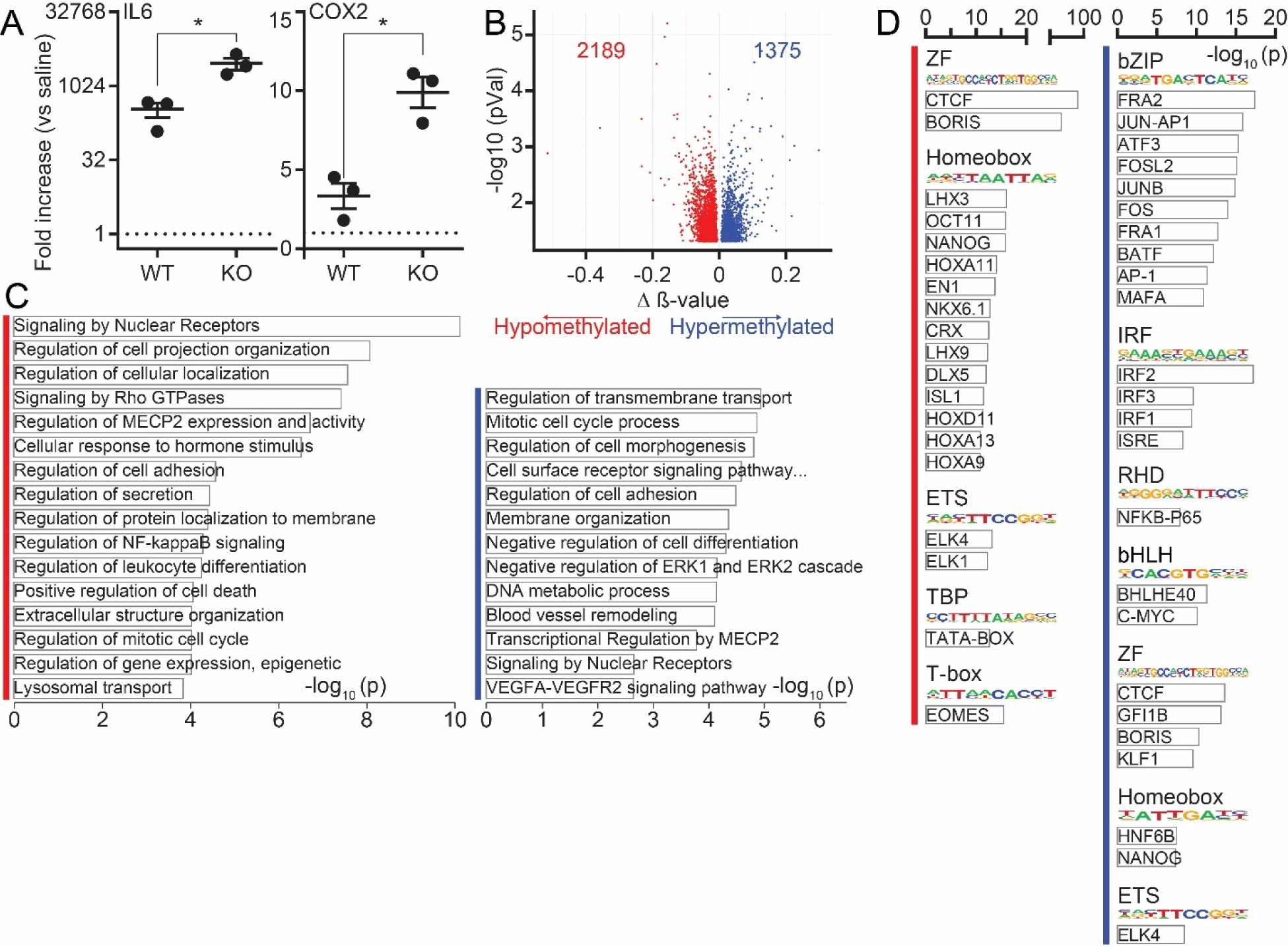
LPS-induced DNA methylation response in the kidney endothelium of mice lacking SOCS3 is associated with the activity of multiple transcription factors. (**A**) RT-qPCR of enriched endothelial cells showing the increased levels of IL6 and COX2 expression in SOCS3iEKO mice after LPS treatment. Mean ± SE fold-change expressed vs WT control mice. (**B**) Volcano plot of differentially methylated positions CpG sites showing the p-value vs. Δβ for LPS-treated WT and SOCS3iEKO mice. The red dots represent significantly hypomethylated CpGs, the blue dots represent significantly hypermethylated CpGs. (**C**) Gene ontology analysis of genes associated with SOCS3iEKO differentially methylated CpG sites showing the most relevant enriched categories (blue mark for the hypermethylated set, and red mark for the hypomethylated set) processed via Metascape (blue mark for the hypermethylated set, and red mark for the hypomethylated set). (**D**) TF motif analysis using HOMER of hyper and hypo methylated gene subsets for LPS-treated WT and SOCS3iEKO mice. Asterisks denote p < 0.05.

### Direct IL-6/STAT3 signaling in vitro promotes sustained DNA methylation and RNA expression changes in HUVEC

The changes in the response of SOCS3iEKO mice suggested a role for IL-6/STAT3 signaling in this context. To directly test this, we challenged cultured HUVEC with a combination of recombinant IL-6 and sIL-6Rα^12, 14, 44, 45^ (herein, IL-6+R, mimicking IL-6 trans-signaling^13^) and obtained genomic DNA and total RNA 6-72 hours post-challenge. Previous work demonstrated strong IL-6+R-induced transcriptional response and barrier function loss within this timeframe^12, 14^. We then performed methylomics assays after genomic DNA bisulfite conversion and RNA-seq from the same samples. No significant changes in DNA methylation were detected 6 or 24 hours post-challenge (not shown). Sustained gene expression changes 72 hours after IL-6+R was demonstrated by RT-qPCR (Figure 3A). Methylomics analysis 72 hours post-IL-6+R identified 431 differentially methylated CpG positions (adjusted p <0.05) (Figure 3B, Supplemental Table 3). Of these, 204 CpG sites were hypomethylated and 227 sites were hypermethylated. GO analysis of the subset of differentially methylated sites located at genes revealed enriched biological processes that closely mimicked those identified in endotoxemic kidney endothelium, including innate immunity, cytokine signaling and cell adhesion, and endothelial cell differentiation (Figure 3C). RNA-seq of the same samples resulted in the IL-6+R-induced upregulation of 316 genes (logFC ≥ 1.5, FDR < 0.05), and downregulation of 195 genes (logFC ≤ -1.5, FDR < 0.05) (Figure 4A, Supplemental Table 4). The over-represented GO categories among the differentially expressed genes were similar to those found in the methylomics analysis, including categories related to cell migration, cell proliferation and response to cytokines (Figure 4B).

**Figure 3.**
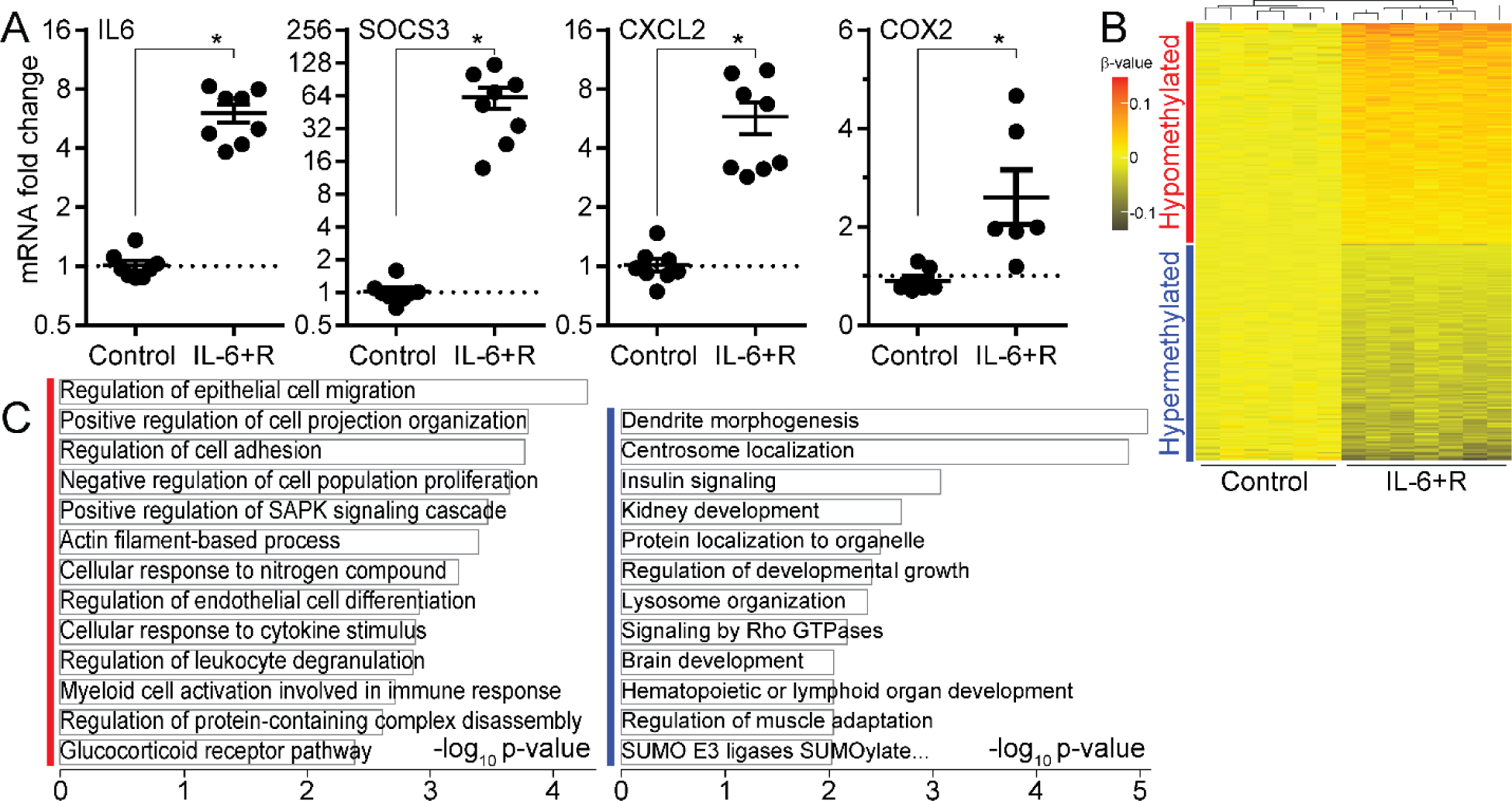
Sustained treatment of HUVEC with IL-6+R induces changes in DNA methylation of inflammatory genes. (**A**) RT-qPCR of HUVEC treated with IL-6 or PBS for 72 hours. Mean ± SE fold change Il-6+R 72 hours vs. control. (**B**) DNA methylation heatmap showing 431 differentially methylated CpG sites between cells treated or not for 72 hours with IL-6+R. The heatmap includes all CpG-containing probes that display significant methylation changes, p-value < 0.05. (**C**) Gene ontology analysis of genes associated with differentially methylated CpG sites showing the most relevant enriched categories, processed via Metascape (blue mark for the hypermethylated set, and red mark for the hypomethylated set).

**Figure 4.**
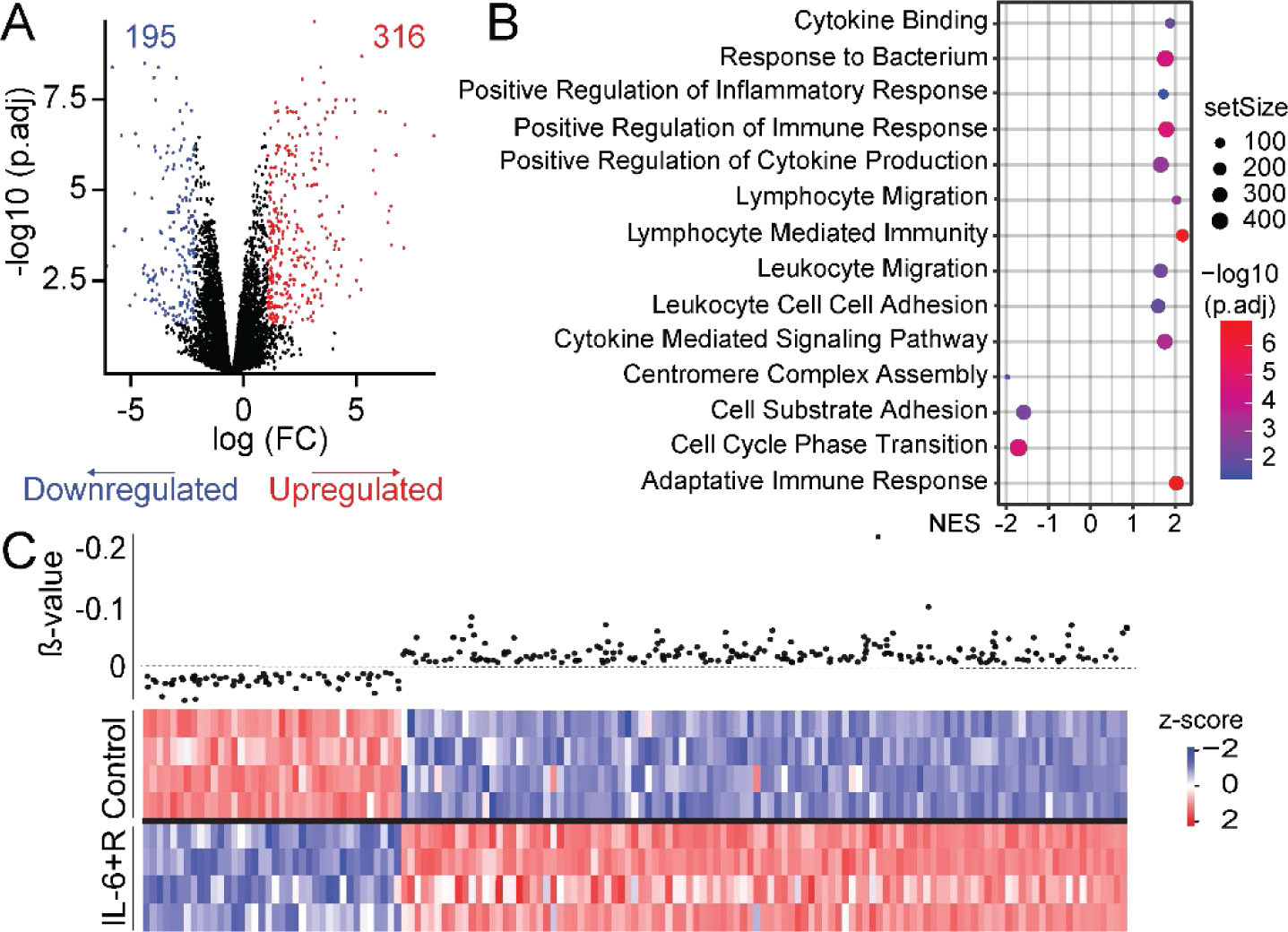
The transcriptional response of HUVEC treated with IL-6+R is associated with changes in DNA methylation. (**A**) Volcano plot. The red dots represent significantly upregulated genes, the blue dots represent significantly downregulated genes (|log2 FC| ≥ 1.5 and adjusted p-value < 0.05), and the black dots represent not significant differentially expressed genes. (**B**) Gene set enrichment analysis (GSEA) of HUVEC treated with IL-6 or PBS for 72 hours, using MSigDB hallmarks (H) as gene sets. Showing normalized enrichment score (NES) (FDR < 0.05). (**C**) Gene expression heatmap (down) and differentially methylated CpG sites (up). Asterisks denote p < 0.05.

As shown in Figure 4C, a comparison between the gene subsets with differential methylation and altered expression in HUVEC showed significant overlap. We identified 145 DEG genes with 269 DMP, comprising 38 genes hypermethylated and downregulated and 107 genes hypomethylated and upregulated (Supplemental table 5). RT-qPCR on selected targets was performed on new biological replicates to confirm this finding (Figure 5A). SERPINA3, NOSTRIN and PLCE1 genes each showed three hypomethylated CpGs in response to an IL-6+R challenge by 72 hours, and consistently, their expression levels were upregulated by 72 hours of IL-6+R treatment. Conversely, TNFSF4 and NAV2 showed three and two hypermethylated CpGs, respectively, and their expression was downregulated by 72 hours of IL-6+R treatment. In all cases, the expression of these genes was not substantially altered by a short IL-6+R treatment, suggesting that the changes observed upon sustained treatment cannot be simply a direct effect of IL-6 transcriptional control (Figure 5A). Notably, the expression changes increased over time and did not reach a maximum until 72 hours of treatment. This behavior is in stark contrast to the fast response of well-known direct STAT3 targets such as SOCS3, COX2 or IL6^12, 14, 15^. To test whether DNA methylation changes can directly alter the expression of these genes, we treated HUVEC for 72 hours with the methyl transferase inhibitor 5-aza-2′-deoxycytidine (5-AZA). This inhibitor led to a significant increase in the expression of SERPINA3, NOSTRIN, PLCE1, and TNSF4, while it did not affect the levels of NAV2 (Figure 5B). STAT3 is the main transcription factor downstream of IL-6. To test whether STAT3 was required for the sustained IL-6-induced change in differentially methylated genes, we performed STAT3 gene knockdown and measured gene expression in cells treated or not with IL-6+R for 72 hours (Figure 5C). STAT3 knockdown abrogated the IL-6+R-induced increasing SERPINA3, NOSTRIN and PLCE1, and the decrease in expression of TNFSF4 and NAV2, demonstrating a causal, albeit probably indirect, role of STAT3 activation in this transcriptional response.

**Figure 5.**
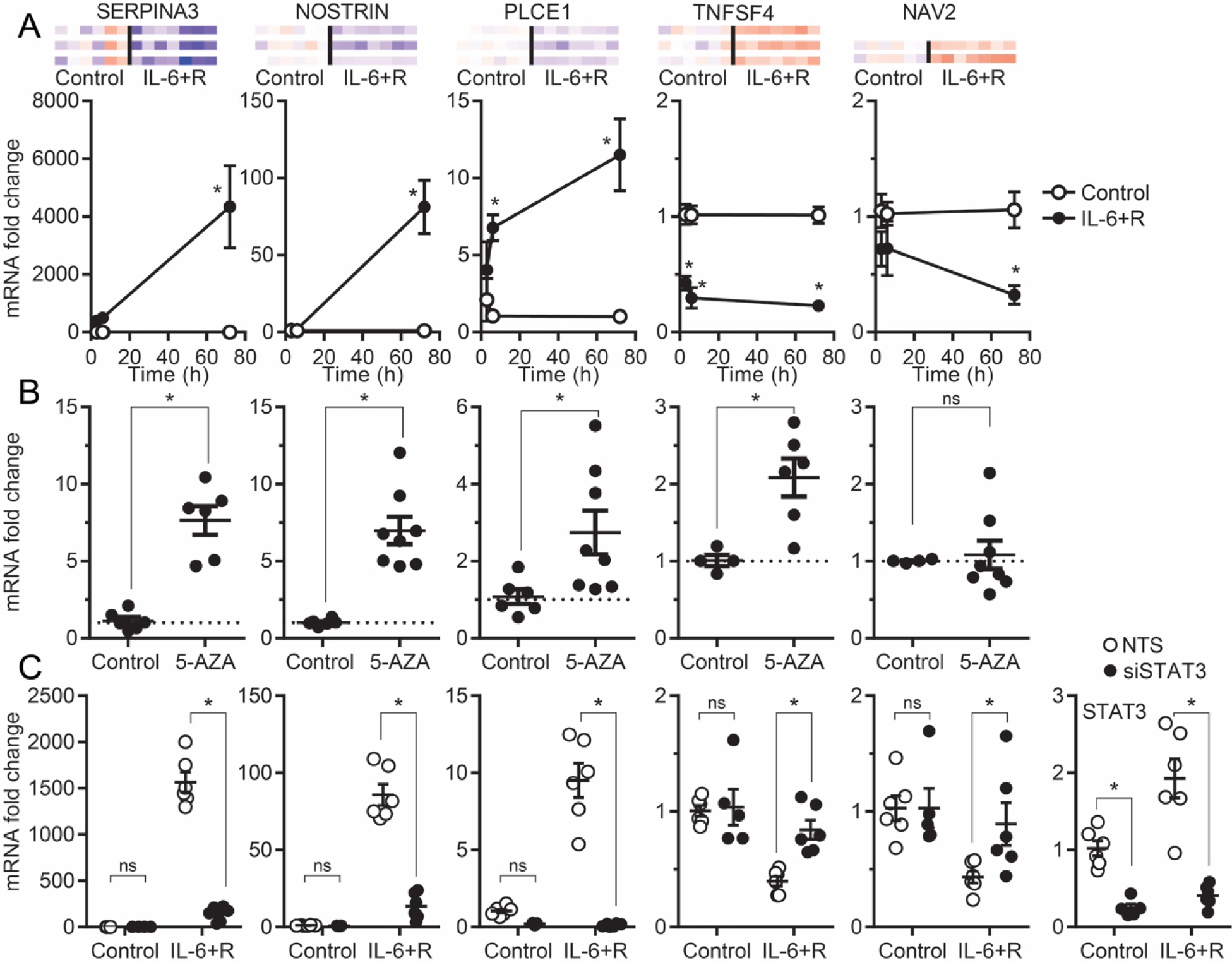
STAT3 activity and DNA methylation regulate gene expression in HUVEC treated for 72 hours with IL-6+R. **A**)DNA hypomethylation on the genes SERPINA3, NOSTRIN and PLCE1 after IL-6+R treatment are associated with increase gene expression and DNA hypermethylated genes TNFSF4 and NAV2 are associated with decrease in the gene expression in HUVEC treated or not with IL-6. Data compiled from at least three independent experiments. Asterisks denote p < 0.05 at 72 hours. (**B**) HUVEC treated with 5-AZA for 72 hours showed an increase gene expression on the genes SERPINA3, NOSTRIN, PLCE1 and TNFSF4. Data compiled from at least three independent experiments. Asterisks denote p < 0.05. (**C**) STAT3 knockdown using siRNA was evaluated by RT-qPCR in HUVEC treated or not with IL-6 with or without siRNA against STAT3. A non-targeting sequence (NTS) siRNA were used as control. Asterisks denote p < 0.05.

### Changes in methylation patterns are retained after IL-6 signaling interruption

We then investigated if the DNA methylation patterns were maintained after IL-6 signaling ended. HUVEC were incubated for 72 hours with IL-6+R or saline as above. Then, cells were either lysed immediately or washed and incubated for an additional 96 hours in regular growth media in the absence of cytokine (Figure 6A). DNA methylation status was assessed by bisulfite conversion and bead arrays as above. Notably, we found that nearly 40% (167 CpGs) of the DMP observed after 72 hours of IL-6 remained altered post-washout (with a mean difference of less than 2%) (Figure 6B, Supplemental Table 6), suggesting that many changes in the endothelial methylome remain in place for prolonged periods, potentially inducing long-lasting transcriptional changes. STAT3 phosphorylation (Figure 6C) and IL-6 mRNA expression (Figure 6D) returned quickly back to control levels after the treatment wash, demonstrating complete removal of the cytokine.

**Figure 6.**
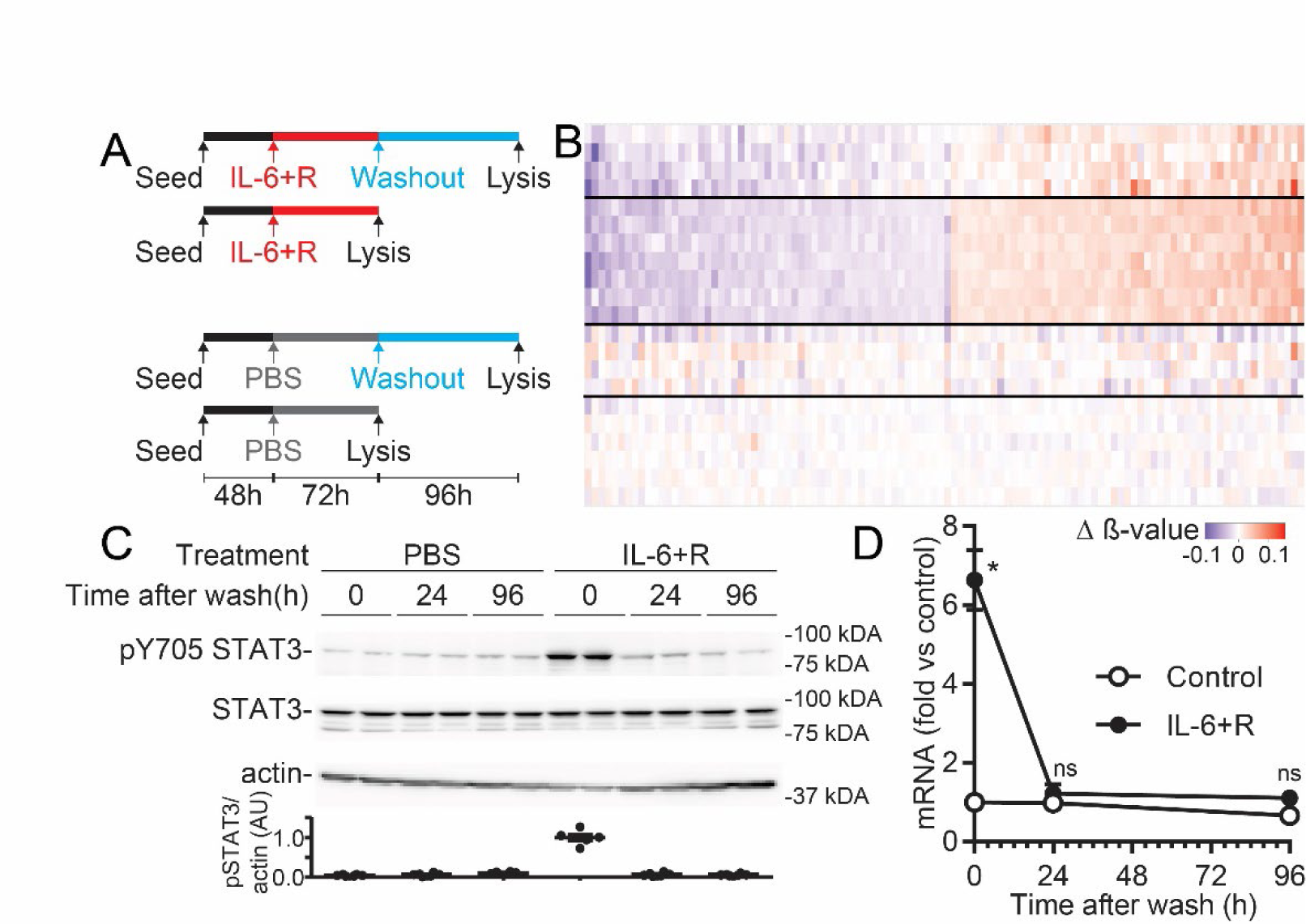
Changes in the endothelial methylome remain in place for prolonged periods in HUVEC. (**A**) Schematic diagram depicting in vitro experiments for IL-6 washout. (**B**) DNA methylation heatmap showing differentially methylated CpGs that are retained after IL-6 washout. (**C**) Cells were treated with the indicated amounts of IL-6+R for 72 hours and wash for 96 hours prior to lysis. Phosphorylated STAT3 and B-actin levels were measured by Western blot. (**D**) Cells were treated as in C prior RNA extraction. IL-6 expression levels were measured by RT-qPCR. GAPDH was used for normalization. Asterisks denote p < 0.05.

### Overlapping TF activities are associated with DNA methylation and gene expression in response to IL-6+R

To gain insights into potential roles for cytokine-induced TF binding, beyond STAT3, to regulatory elements driving the changes in the endothelial DNA methylation, we performed analysis of TF motif enrichment of the IL-6+R differentially methylated gene set (Figure 7A). Notably, the motif enrichment closely mimics the findings obtained previously from endotoxemic kidney endothelium. Motifs enriched in hypomethylated genes included those binding STAT1 and STAT3, as well as ETS and homeobox families; while those enriched in hypermethylated genes included those binding AP1 family members (including JunB, cFos and BATF) and IRFs. Our results also show enrichment in AP1 binding sites near hypermethylated positions that remained differentially methylated 96 hours after IL-6 wash (Figure 7B), strongly arguing for a sustained epigenetic response to the activation of these transcription factors.

**Figure 7.**
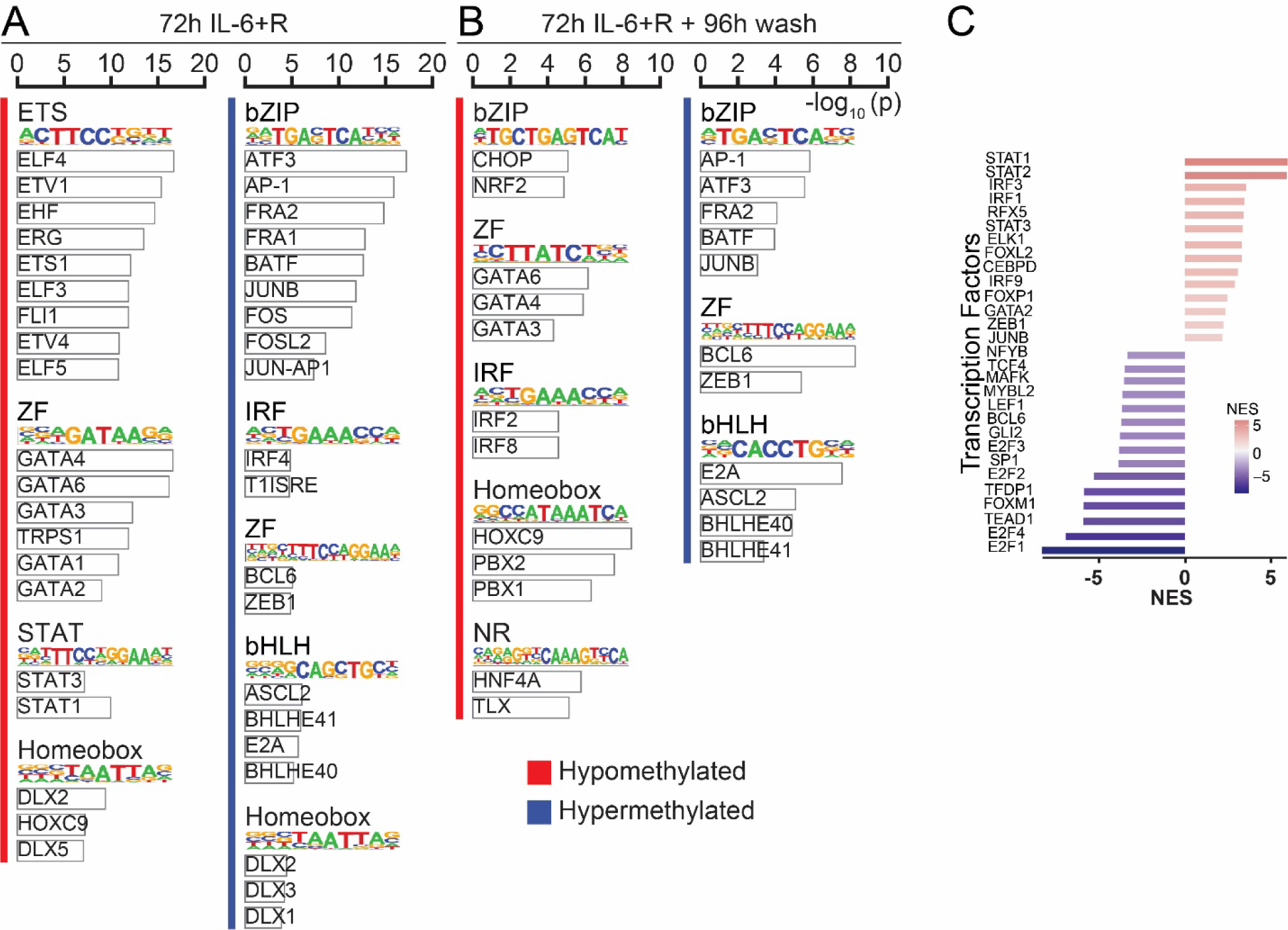
Multiple transcription factors are enriched in the gene sets that are differentially methylated or differentially expressed upon sustained IL-6+R signaling. (**A**) TF motif analysis of hyper and hypo methylated gene subsets between cells treated or not for 72 hours with IL-6+R. TF family and factor (selected TF with p ≤ 1e−05 for hypermethylated and hypomethylation regions) Motif logo is representative of the TF family. (**B**) TF motif analysis of differentially methylated CpG between IL-6 for 72 hours and 96 hours wash. TF family and factor (selected TF with p ≤ 1e−05 for hypermethylated and hypomethylation regions) Motif logo is representative of the TF family. (**C**) Selected TF activities inferred with DoRothEA from gene expression in HUVEC treated with IL-6 for 72 hours. Showing normalized enrichment score (NES).

We then performed discriminant regulon expression analysis of the HUVEC RNA-Seq data to test if these TF were associated with gene expression changes (Figure 7C). Increased STAT activities further confirmed the sustained IL-6 signaling 72 hours post-challenge. Other TF identified closely overlapped with the motifs associated with differentially methylated genes, including the AP1 member JunB and several IRFs.

Consistent with a potential role for JunB in these responses, we found that JunB mRNA (Figure 8A) and protein (Figure 8B) levels quickly increase upon an IL-6+R treatment. Given its constitutively nuclear localization in HUVEC, it is likely that protein levels are the main mechanism of regulation for this transcription factor (Figure 8C). We then performed knockdowns to test a causal role for JunB in IL-6+R responses. As shown in Figure 8D, siRNAs against JunB led to an ∼90% decrease in protein expression. COX2 and PCDH17 were identified as potential JunB targets, according to the ChIP-X enrichment analysis (CHEA) database^46^. As shown in Figure 8E, JunB knockdown in HUVEC abrogated IL-6-induced increase in COX2 and PCDH17 expression. In contrast, this knockdown did not prevent the increase in expression of SOCS3, a direct STAT3 target gene (Figure 8F).

**Figure 8.**
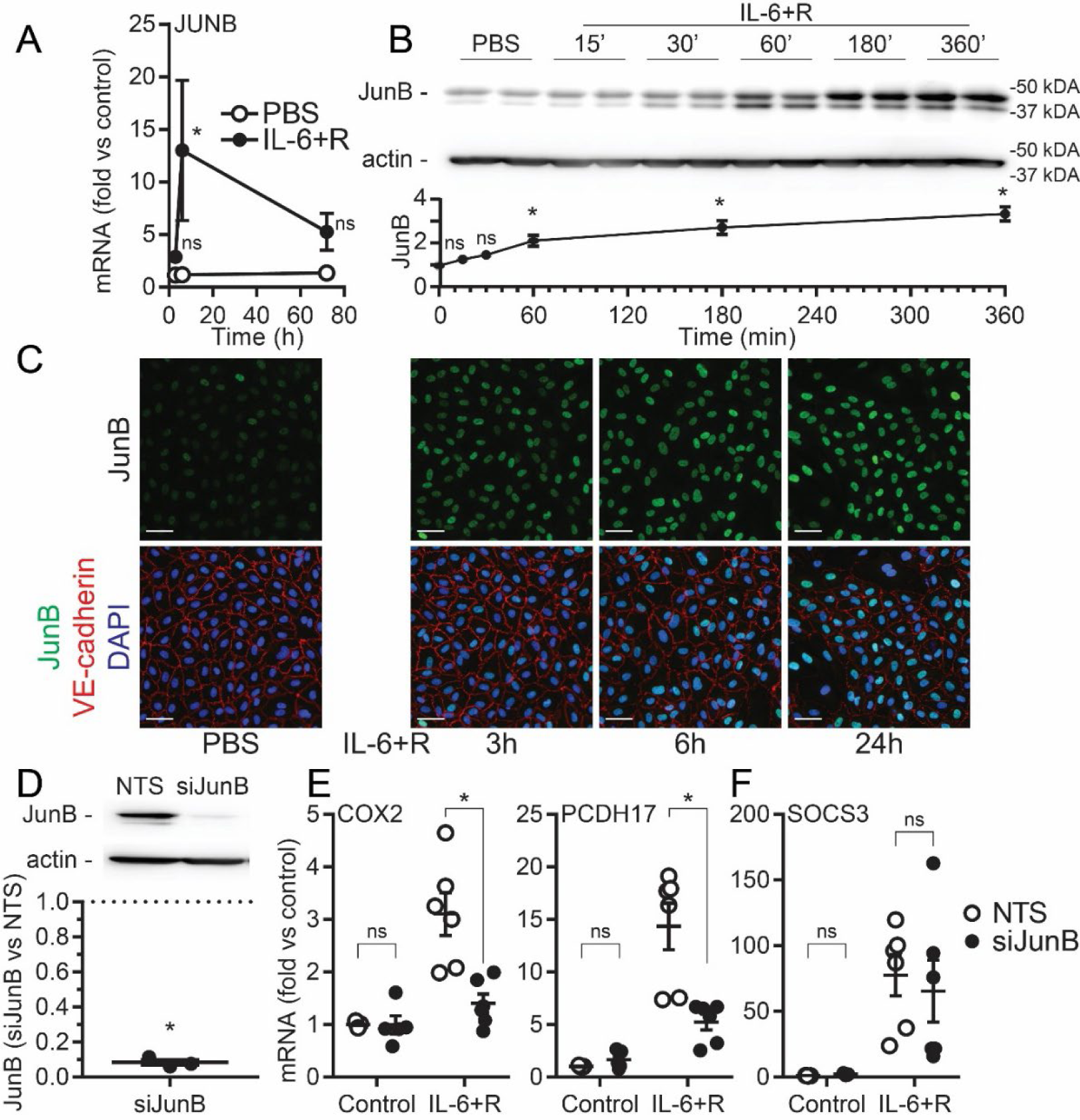
JunB is required for IL-6+R-induced gene expression changes. (**A**) JunB RT-qPCR of HUVEC treated with IL-6 or PBS. Mean ± SE fold change Il-6+R vs. control. (**B**) Cells were treated with the indicated amounts of IL-6+R prior to lysis. JunB levels were measured by Western blot (**C**) Immunofluorescent Staining of Confluent Human Umbilical Vein Endothelial Cells (HUVEC) treated with IL-6 and soluble IL-6Rɑ (IL-6+R) as an inducer of acute inflammation vs. PBS Control. HUVEC were treated on 3 HR, 6 HR, and 24 HR timepoints. Following treatment, cells were fixed, stained with anti-JunB (Green), anti-VE-cadherin (red), and DAPI (blue). (**D**) JunB knockdown using siRNA was evaluated by WB. A non-targeting sequence (NTS) siRNA were used as control. (**E**) Gene expression for SOCS3, COX2 and PCDH17 in HUVEC treated or not with IL-6 for 72 hours with or without siRNA against JunB. Asterisks denote p < 0.05.

### AP1 activity is associated with the endothelial transcriptional response in endotoxemic kidneys

We then performed translating ribosome affinity purification (TRAP) RNA sequencing (TRAP-Seq) from kidneys derived from LPS-treated control or SOCS3iEKO mice. Data from WT and SOCS3iEKO kidney TRAP-Seq shows that while loss of SOCS3 does not lead to significant changes in the endothelial translatome of otherwise healthy mice, it greatly affects the gene expression signature upon an LPS challenge (Figure 9A). Similar to our findings using magnetic bead sorting, we found that SOCS3 depletion exacerbated the LPS-induced increase in IL-6 and COX2 expression (Supplemental table 7). We observed a strong transcriptional response to LPS (565 genes were significantly upregulated, while 2688 genes were downregulated), demonstrating a broad endothelial response to systemic inflammation (Figure 9B). Notably, many of the genes highly induced by LPS >2-fold in SOCS3iEKO but not in WT mice include those we previously found associated with an acute response to IL-6 in HUVEC^12^, including LIPG, RHOU, DDX58, F3, BATF3, and EPSTI1 (Supplemental table 7). Genes induced by LPS in WT mice but that SOCS3iEKO mice had at least a further 2-fold increase vs WT include PTGS2 (coding for COX2), CXCL10, CEBPD, MX1 and IL6 itself, genes that were also found previously as quickly induced by IL-6+R in HUVEC. Metascape analysis of this data again showed enrichment in processes such as the inflammatory response and cell cycle regulation (Figure 9C). Notably, motif enrichment analysis demonstrated increased activity of not only STAT1 and STAT3, but also AP1 members BATF, cFos, and JunB and IRF (Figure 9D).

**Figure 9.**
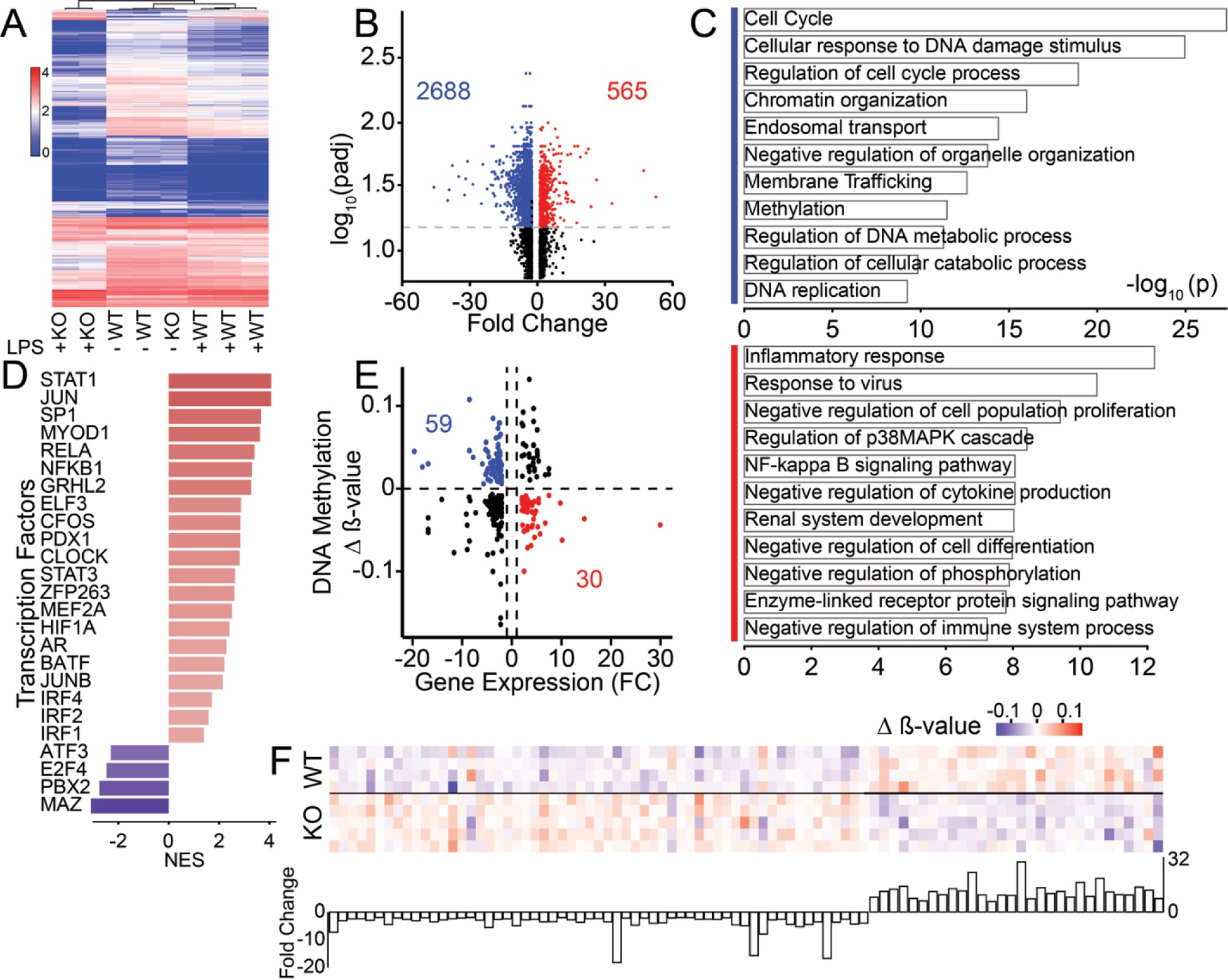
Loss of SOCS3 in the kidney endothelium leads to altered gene expression that is associated with transcription factor activation and DNA methylation. (**A**) Heatmap and unbiased clustering from all genes induced or inhibited >1.5-fold by LPS (4842 genes, log scale of expression levels). (**B**) Volcano plot. The red dots represent significantly upregulated genes, the blue dots represent significantly downregulated genes (|log2 FC| ≥ 2 and FDR < 0.05), and the black dots represent insignificant differentially expressed genes. (**C**) Gene ontology analysis of genes associated with SOCS3iEKO DEG showing the most relevant enriched categories (blue mark for the downregulated genes, and red mark for the upregulated genes). (**D**) Selected TF activities inferred with DoRothEA from gene expression in SOCS3iEKO kidney endothelium mice after LPS treatment. (**E**) Correlation analysis between gene expression and DNA methylation changes. The x-axis is the log2 fold change of gene expression between LPS-treated WT and SOCS3iEKO mice. The y-axis is the delta beta of DNA methylation change of mapped genes. (**F**) Correlation between DNA methylation changes (heatmap) and mRNA expression (barplot).

Similar to our findings in HUVEC, the gene subsets with differential methylation and altered expression in endotoxemic kidneys showed significant overlap. As shown in Figure 9E, 89 genes were identified with both DNA methylation and gene expression variations (Supplemental table 8), comprising 60 hypermethylated positions and downregulated expression and 37 hypomethylated positions and upregulated expression (Figure 9F). Consistent with a role for STAT3, IRF and AP-1 promoting gene expression, many genes associated with STAT3 binding showed decreased methylation and increased expression (Supplemental table 8).

## Discussion

Here, we provide evidence for an IL-6-mediated coordinated response in endothelial cells that comprises both epigenetic and transcriptional changes. We show that acute transcription through AP1, STAT, and IRF families is associated with DNA methylation changes in multiple pro-inflammatory genes that suggests a coordinated response to acute events with the potential of modulating long-term vascular consequences. This bi-directional response, in which not only DNA methylation affects gene expression, but also acute transcriptional events lead to epigenetic marks, could explain at least in part the chronic sequelae after shock. Furthermore, our in vitro experiments after IL-6 removal suggest that this cytokine may elicit transcriptional changes long after the resolution of inflammation.

Endothelial dysfunction is a common feature of many inflammatory diseases^2^. Septic shock survivors have increased risk of developing long-term complications, including endothelial dysfunction, which can have significant health consequences^47–49^. However, little is known about the regulation within the endothelium of injured organs of specific transcription factor networks and epigenetic responses during and after resolution of inflammation. A better understanding of these mechanisms is key for designing therapies for reducing the incidence of endothelial dysfunction and its complications in septic shock survivors. DNA methylation modifications in septic patients are predominantly examined in mixed populations of circulating cells^16, 18, 19^. One study identified 668 methylation sites that were altered between patients with sepsis from those with non-septic critical illness^16^. Indeed, across functional enrichment analysis, DNA methylation alterations have been identified in methyltransferase activity, cell adhesion, and cell junctions^16^. Furthermore, several studies showed that the expression of enzymes involved in epigenetic modifications and chromatin remodeling varies with severity and cell type^18, 50, 51^. In a recent study, it was found that alterations in DNA methylation in human monocytes following a septic episode correlate with circulating IL-10 and IL-6 levels, suggesting a potential mechanism downstream involving the generation of defective DNA methylation alterations^52^. Notably, this study also showed significant changes in the methylation status of IRF and STAT response genes^52^. A second study showed that LPS leads to changes in monocyte DNA methylation that are concomitant with the upregulation of inflammatory-related genes, and these involve the JAK2/STAT pathway^53^. In addition, a mouse study identified 200 promoter genes with differentially methylated promoters in whole kidneys 24 hours after ischemia and reperfusion injury, of which 79 maintained the difference 7 days after reperfusion^54^. Another report showed 1,721 genes are differentially methylated in rat lung tissue in response to LPS-induced acute lung injury^55^. Together, these studies argue for an important role for DNA methylation in response to injury, but the cell-type specific changes and their mechanisms within the failing organs remain unknown. Our innovative approaches of enriching endothelial cells from failing kidneys for methylome analyses and obtaining the endothelial translatome through a translating ribosome affinity purification strategy allowed us to directly assess the response to an acute inflammatory challenge within the kidney endothelium.

We identified multiple modifications in the DNA methylation on the kidney endothelium in the endotoxemic mice that were associated with an increased inflammatory response. We focused on the kidney because of its well characterized injury in mouse models of systemic inflammation^12, 56^ and because we previously showed a dramatic loss of kidney function in endotoxemic SOCS3iEKO mice^12^. Consistently, loss of endothelial SOCS3 led to a differential epigenetic signature in response to LPS compared to WT mice. A gene ontology analysis showed an enrichment of pathways downstream of inflammatory cytokines and cell adhesion, consistent with an acute, severe inflammatory reaction. Also, the gene ontology analysis showed an enrichment of pathways associated with chromatin organization and regulation of epigenetics. Notably, IL-6 treatment on cultured endothelial cells led to differential methylation of a similar proinflammatory and epigenetic regulation gene set. This specific epigenetic response, consistent with the transcriptional changes induced by LPS and IL-6, led us to investigate a potential mechanism to coordinate these two responses. Bioinformatics analysis of the SOCS3iEKO kidney epigenetic data suggested an enrichment of binding sites for several families of transcription factors, most notably AP1 and IRF. A strikingly similar enrichment was observed in IL-6-treated HUVEC, consistent with a key role for IL-6 in the endothelial response to IL-6^12, 15^. Bioinformatic analysis of the endothelial transcriptional response both in vivo and in vitro again suggested an increased activity of AP1 and IRF transcription factor families, together with the expected STAT3 activity^12, 14, 57^. Here, we confirmed that STAT3 and DNA methylation both regulate the levels of expression of several differentially methylated genes, suggesting a novel role for STAT3 not only in regulating gene expression, but also in enabling epigenetic changes at these loci.

The expression of many AP1 members, including JunB, cFos and BATF, is quickly induced by an IL-6 challenge in HUVEC and is strongly induced in the endotoxemic kidney endothelium. AP1 transcription factors are involved in activating genes involved in the inflammatory response, including cytokines, chemokines, and adhesion molecules^58–62^, and AP1 inhibitors may reduce inflammation in various disease models^59, 63^. Our study observed enrichment in AP-1 binding sites nearby hypermethylated positions 72 hours after an IL-6+R treatment that remained differentially methylated 96 hours after wash. To begin assessing a causal role for these transcription factors, we knocked down JunB in HUVEC. Notably, the increased expression of COX2 was abolished in cells depleted of this transcription factor. Many other genes associated with AP1 binding, however, did not. This suggests a strong specificity for AP1 members in the regulation of different AP1-reponsive genes. Alternatively, gene compensation from other AP1 members may have masked any JunB role. The specific role of JunB in regulating the immune response can vary depending on the specific cell type, stage of development, and the presence of other regulatory elements^64–66^. The exact mechanism by which JunB regulates COX-2 expression is not well understood^67, 68^. Further work is required to fully understand the roles for AP1 members in this context.

In conclusion, we demonstrate that the epigenetic changes in the mouse kidney endothelium in response to endotoxemia are dependent on the expression of the IL-6 negative regulator SOCS3. In vitro, we show that IL-6 signaling directly promotes a sustained alteration in DNA methylation that may mediate many of the changes in endothelial gene expression. Our cross-omics analyses of transcriptome and DNA methylome analysis suggest the potential involvement of specific transcription factors in the regulation of these DNA methylation changes, providing potential candidate targets to limit the long-term damage induced by shock. This study highlights the importance of epigenetic modifications in endothelial function and provides a foundation for developing novel therapeutic interventions for inflammatory diseases.

## Supplemental table and figures

Supplemental Table 1. Differentially methylated positions in mice treated or not with LPS

Supplemental Table 2. Differentially methylated positions in LPS-treated mice lacking or not SOCS3 in the endothelium

Supplemental Table 3. Differentially methylated positions in HUVEC 72h after IL-6+R treatment

Supplemental Table 4. Differentially expressed genes in HUVEC 72h after IL-6+R treatment

Supplemental Table 5. Correlation of Differentially methylated positions and differentially expressed genes in HUVEC 72h after IL-6+R treatment

Supplemental Table 6. Differentially methylated positions in IL-6+R-treated HUVEC after a 96-hour wash

Supplemental Table 7. Differentially expressed genes in the kidney endothelium of WT and SOCS3IEKO mice treated or not with LPS

Supplemental Table 8. Correlation of Differentially methylated positions and differentially expressed genes in the kidney endothelium of WT and SOCS3IEKO mice treated or not with LPS

Supplemental Table 9. Vendor, catalog numbers, and concentrations for reagents

Supplemental Table 10. Primers sequences

Supplemental Table 11. Antibodies, including vendor catalog numbers, RRIDs, and the specific concentrations and blocking solutions

