## Supplemental Table 11 for "Shock drives a highly coordinated transcriptional and DNA methylation response in the endothelium"

**Supplemental Table 11 – List of antibodies**

| **For Western blotting** | | | | | | | | |
| --- | --- | --- | --- | --- | --- | --- | --- | --- |
| **Target** | **Species** | **Clone** | **Conjugate** | **Vendor** | **Catalog No** | **RRID** | **Dilution** | **Blocking solution** |
| STAT3 | rabbit | D1A5 |  | Cell Signaling | 8768S | AB_2722529 | 1:5000 | BSA, 0.1%, 0.1%  Tween in PBS |
| pY705-STAT3 | rabbit | D3A7 |  | Cell Signaling | 9145L | AB_2491009 | 1:5000 | BSA, 0.1%, 0.1%  Tween in PBS |
| JunB | rabbit | C37F9 |  | Cell Signaling | 3753 | AB_2130002 | 1:1000 | BSA, 0.1%, 0.1%  Tween in PBS |
| β-actin | mouse | AC-15 |  | Millipore Sigma | A5441 | AB_476744 | 1:10000 | 5%BSA, 0.1%  Tween in PBS |
| Peroxidase  AffiniPure Goat  Anti -Mouse IgG  (H+L) | goat | polyclonal | HRP | Jackson IR | 115-035-062 | AB_2338504 | 1:5000 | Same as primary  Ab blocker |
| Peroxidase  AffiniPure Goat  Anti -Rabbit IgG  (H+L) | rabbit | polyclonal | HRP | Jackson IR | 111-035-003 | AB_2313567 | 1:5000 | Same as primary  Ab blocker |

| **For Immunofluorescence** | | | | | | | | |
| --- | --- | --- | --- | --- | --- | --- | --- | --- |
| **Target** | **Species** | **Clone** | **Conjugate** | **Vendor** | **Catalog No** | **RRID** | **Dilution** | **Blocking solution** |
| JunB | rabbit | C37F9 |  | Cell Signaling | 3753 | AB_2130002 | 1:1000 | 5% FBS in PBS-TX |
| VE-cadherin | goat | AC-15 |  | R&D Systems | AF938 | AB_355726 | 1:100 | 5% FBS in PBS-TX |
| Anti -Rabbit IgG | donkey | polyclonal | Alexa Fluor 647 | Invitrogen | A-31573 | AB_2536183 | 1:500 | 5% FBS in PBS-TX |
| Anti- Goat IgG | donkey | polyclonal | Alexa Fluor 594 | Invitrogen | A-11058 | AB_2534105 | 1:500 | 5% FBS in PBS-TX |

| **For EC enrichment** | | | | | | | | |
| --- | --- | --- | --- | --- | --- | --- | --- | --- |
| **Target** | **Species** | **Clone** | **Conjugate** | **Vendor** | **Catalog No** | **RRID** | **Dilution** | **Blocking solution** |
| CD326 (Ep-CAM) | rat | G8.8 | Biotin | BioLegend | 118204 | AB_1134174 | 0.5mg/mL |  |
| CD31 | rat | MEC13.3 | Biotin | BioLegend | 102504 | AB_312910 | 0.5mg/mL |  |
| CD45 | rat | 30-F11 | Biotin | BioLegend | 103104 | AB_312969 | 0.5mg/mL |  |
