## Supplemental Table 9 for "Shock drives a highly coordinated transcriptional and DNA methylation response in the endothelium"

| **M&M section** | **Reagent or disposable** | **Vendor** | **Catalog Number** | **Concentration/Dose** |
| --- | --- | --- | --- | --- |
| **Mice** | Tamoxifen | Sigma Aldrich | T5648 | 2 mg in 100 μL |
|  | Peanut Oil | Sigma Aldrich | P2144 | N/A |
|  | Lipopolysaccharides | Sigma Aldrich | L4391 | 250 μg/ 250 μL |
| **Endothelial Enrichment** | Collagenase Type 1 | Worthington Biochemicals | LS004196 | 0.2% |
|  | RNAse A | Sigma Aldrich | E866 | 100 mg/ml |
|  | Dispase II | Roche | 10295825001 | 0.2% |
|  | Dneasy Blood and Tissue kit | Qiagen | 69504 | As per instructions |
|  | RNEasy Plus Micro Kit | Qiagen | 74034 | As per instructions |
|  | BD Falcon cell strainer 70um pore size | Thermo Fisher | 352350 | N/A |
|  | Dynabeads M-280 Streptavidin | Thermo Scientific | 11205D | 1mg/100 μL |
|  | Trypsin-EDTA Solution 1X | Millipore Sigma | 54917C-100ML | 1X |
| **Cell Culture** | Recombinant Human IL6 protein | R&D Systems | 206-IL-200/CF | 200 ng/mL |
|  | Recombinant human IL6r alpha, CF | R&D Systems | 227-SR-025/CF | 100 ng/mL |
|  | phenol red-free EBM 2 media | PromoCell | C-22211 | N/A |
|  | Penicillin-Streptomycin Solution, 100x | Corning | 30-002-CI-PK | 1X |
|  | Gelatin | Millipore Sigma | ES-006-B | 0.1% |
|  | DAPI (4',6-Diamidino-2-Phenylindole) | Thermo Fisher Scientific | D3571 | 1 ug/ml |
|  | Fluoroshield with 1,4-Diazabicyclo [2.2.2]  octane | Millipore Sigma | F937 |  |
|  | 5-Aza-2′-deoxycytidine | Millipore Sigma | A3656 | 5 µM |
| **Gel Electrophoresis and**  **Immunoblotting** | Complete protease inhibitor mixture | Roche Applied | 11697498001 | 1X |
|  | PhosSTOP phosphatase inhibitor mixture | Roche Applied | 4906845001 | 1X |
|  | Sodium fluoride | Millipore Sigma | S-1504 | 100 mM |
|  | Phenyl arsine oxide | Millipore Sigma | P-3075 | 100 µM |
|  | Sodium pyrophosphate decahydrate | Millipore Sigma | 221368-100G | 10 mM |
|  | Sodium orthovanadate | Millipore Sigma | S6508 | 100 µM |
|  | Transblot Turbo RTA Mini Nitrocellulose - Transfer kit | Bio-Rad | 1704270 | N/A |
|  | Clarity Western ECL Substrate | Bio-Rad | 1705061 | As per instructions |
|  | Bio-Rad Clarity Max Western ECL Substrate | Bio-Rad | 1705062 | As per instructions |
| **RNA isolation and RTqPCR** | PrimeScript RT Master Mix | Clontech | RR036B | 1X |
|  | TRIzol reagent | Invitrogen | 15596018 | 1X |
|  | iTaq Univer SYBR Green Supermix | Bio Rad | 1725125 | 1X |
| **siRNA** | lipofectamine RNAiMAX transfection reagent | Invitrogen | 13778150 | 6 pmol siRNA/μl lipid |
|  | ON-TARGETplus non-targeting control pool | Horizon Discovery | D-001810-10-20 | 50 nM |
|  | Opti-MEM | Thermo Fisher Scientific | 31985070 | 1X |
|  | ON-TARGETplus 1 set JunB siRNA | Horizon Discovery | J-003269-09 | 50 nM |
|  | ON-TARGETplus 1 set STAT3 siRNA | Horizon Discovery | J-003544-07 | 50 nM |
| **TRAP** | Avanti Polar Lipids, 1,2-diheptanlyl-sn-glycerol | Avanti Polar Lipids | NC9999043 | 300mM |
|  | RNAsin Plus | Promega | PAN2615 | 40 U/ml |
|  | Protein G beads | Thermo Fisher/Invitrogen | 10004D |  |
|  | Ab for TRAP | Memorial Sloan Kettering CC | HTZGFP-19C8 |  |
|  | Ab for TRAP | Memorial Sloan Kettering CC | HTZGFP-19F7 |  |
|  | HBSS no calcium, mg, phenol red | Gibco | 14175079 | 1X |
|  | Complete mini edta free easy pack | Roche | 45-4693159001-ea |  |
|  | PhosStop 20 tablets | Roche | 45-4906837001-ea |  |
|  | Cyclohexamide 1 g | Sigma | 45-c1988-1g-ea | 100 µg/m |
|  | DTT-biotech grade | VWR Life Sciences | 97061-340 | 0.5 mM |
|  | Potassium Chloride ACS | VWR Life Sciences | 97061-566 | 350 mM |
|  | HEPES Free Biotechnology grade | Sigma | 45-H4034-100G-EA | 1M |
|  | Magnesium Chloride | Honeywell | 27678-63020-1L-EA | 1M |
|  | Diethyl pyrocarbonate (DEPC), 5ml | Sigma | 45-D5758-5ML-EA |  |
|  | Sodium Bicarbonate | Fisher Scientific | S233 | 4 mM |
|  | (D) + Glucose | Acros | 41095 |  |
|  | Igepal | MP Biomedical | CA-630 | 1X |
|  | Disposable pellet pestles with 1.5ml microtube | DWK Kimble Kontes | 749520-0000-cs |  |
|  | Cordless Pestle Motor | VWR Life Sciences | 47747-370 |  |
|  | Diamond Midi Centrifuge Tube, 5.0mL, PP, Separate Red Screw Cap, Graduated | Globe Scientific | 111580 |  |
|  | RNeasy Plus Micro Kit | Qiagen | 74034 |  |
