## Supplemental Table 10 for "Shock drives a highly coordinated transcriptional and DNA methylation response in the endothelium"

**Mouse RT-qPCR primer sequences**

| **Primer name** | **Forward Sequence** | **Reverse Sequence** |
| --- | --- | --- |
| CDH1 | GGTCATCAGTGTGCTCACCTCT | GCTGTTGTGCTCAAGCCTTCAC |
| COX2 | GCGACATACTCAAGCAGGAGCA | AGTGGTAACCGCTCAGGTGTTG |
| IL-6 | TACCACTTCACAAGTCGGAGGC | CTGCAAGTGCATCATCGTTGTTC |
| VWF | AACAGACGATGGTGGACTCAGC | AACAGACGATGGTGGACTCAGC |

**Human RT-qPCR primer sequences**

| **Primer name** | **Forward Sequence** | **Reverse Sequence** |
| --- | --- | --- |
| ADAM19 | CGAGAAGGTGAATGTGGCAGGA | AGCTCTGACACTGGATCTTCCC |
| B-actin | CACCATTGGCAATGAGCGGTTC | AGGTCTTTGCGGATGTCCACGT |
| COX2 | CGGTGAAACTCTGGCTAGACAG | GCAAACCGTAGATGCTCAGGGA |
| CXCL2 | GGCAGAAAGCTTGTCTCAACCC | CTCCTTCAGGAACAGCCACCAA |
| HSD17B | TCCAACCTGGAGGCTTCCTAAC | GCTGTGCTAAGATGTAGTCCTGG |
| IL-6 | TACCACTTCACAAGTCGGAGGC | CTGCAAGTGCATCATCGTTGTTC |
| LAMP3 | TGGGAGCCTATTTGACCGTCTC | GCTGACAACTGGAGGCTCTGTT |
| NAV2 | TGGAGCCAAGTACCCAGATGTG | GAGGATTGGAGATGACCACCGA |
| NOSTRIN | AGACCTCAACCCAGCCATCCTT | GAAGAACCTGGATTGCTCTGCC |
| PCDH17 | CAAAGGCTCCTGCTGTGACATG | GACCAAGCACTCGGCATTCATC |
| PLCE1 | CCTGGGCATAAGCACTACCAAG | GTCTTGAGGATCAGAACCACTCC |
| RHOU | ACTGCCTTCGACAACTTCTCCG | GAGCAGGAAGATGTCTGTGTTGG |
| SERPINA3 | CCTGAACGACATACTTCTCCAGC | CATCAAGCACAGCCTTATGGACC |
| SOCS3 | CATCTCTGTCGGAAGACCGTCA | GCATCGTACTGGTCCAGGAACT |
| TNFSF4 | CCTACATCTGCCTGCACTTCTC | TGATGACTGAGTTGTTCTGCACC |
